# Para-cellular and transcellular diapedesis are divergent but inter-connected evolutionary events

**DOI:** 10.1101/2020.09.21.307066

**Authors:** Norwin Kubick, Pavel Klimovich, Patrick Henckel, Michel-Edwar Mickael

## Abstract

Infiltration of the endothelial layer of the blood-brain barrier by leukocytes plays a critical role in health and disease. When passing through the endothelial layer during the diapedesis process lymphocytes can either follow a para-cellular route or a transcellular one. There is a debate whether these two processes constitute one mechanism, or they form two evolutionary distinct migration pathways. We used phylogenetic analysis, HH search, ancestor sequence reconstruction together with functional specificity and positive selection analysis to investigate this intriguing question further. We found that the two systems share several ancient components, such as RhoA protein that plays an important role in controlling actin movement in both mechanisms. However, some of the key components differ between these two transmigration processes. CAV1 genes emerged during Trichoplax adhaerens and it was only reported in trans-cellular process. Para-cellular process core proteins had at least two distinct starting points. First, during drosophila emergence, Tre1 which is homologous to melatonin GPCR receptor diverged. Secondly, PECAM1 emerged from FASL5/3 during elephant shark divergence. Lastly, both systems employ late divergent genes such as ICAM1 and PECAM1. Taken together our results suggest that these two systems constitute different yet interconnected mechanisms of immune cells infiltrations of the brain. Our analysis indicates that this system coevolved with immune cells, evolving to a higher level of complexity in association with the evolution of the adaptive immune system.

## Introduction

### Introduction of blood brain barrier infiltration

The regulation of lymphocytes infiltration through the endothelial cells of the BBB is an intriguing phenomenon which plays a vital role in homeostasis and in disease (Daneman & Prat, 2015). In homeostasis, high BBB integrity and lack of lymphocytes adhesion molecules reduce CD4+ T cells infiltration to the brain. The BBB integrity is assured by three factors(Kubick, Flournoy, Enciu, Manda, & Mickael, 2020). The first is the existence of tight junctions which are controlled by ZO1, ZO2 and ZO3 as well as occludin(Ronaldson & Davis, 2011). The second factor is the junctional adhesion molecules and there are three of them (JAM1, JAM2 and JAM3) in addition to ESAM(Derada Troletti, de Goede, Kamermans, & de Vries, 2016). The third one is the existence of cadherins which represent the adhesion junctions between adjacent endothelial cells (Derada Troletti et al., 2016). The main protein that controls this process is VE-Cadherin and it is connected to the actin of the cell through alpha and beta catenin(Banks, 2012). This compact system insures that CD4+ T cells access to the brain is limited during hemostasis. However, during pathogens invasions, to the brain, the endothelial cells confirmation changes to increase the probability of successful infiltrations of the CD4+ T cells to the brain. A significant downregulation of the junctional proteins such as occludin, JAMs and ZO1,ZO2 and ZO3 has been reported. Additionally, an upregulation in leucocytes adhesion molecules such as VCAM and ICAM as well as activation signals such as CCL9 was observed(Banks, 2012). The main function of these molecules is to capture, activate and provide tight adhesion to the migrating CD4+ T cells into the brain, in what is known as diapedesis.

Diapedesis can take one of two forms. The first is known as para-cellular transmigration and the second is known as transcellular transmigration(Muller, 2011). In para-cellular infiltration several proteins interact together in order to widen the space between endothelial cells. Mainly, ICAM 1 and VCAM1 already connected with ligands on CD4+ T cells during the capture and the adhesion phases become activated and in turn activate the RhoA, Src pathway, that phosphorylates the β-catenin thus freeing VE-cadherin(Adam, 2015). Following that, LBRC (lateral border recycling Compartment) stores the junctional proteins(Muller, 2015; Wettschureck, Strilic, & Offermanns, 2019). PECAM1 expressed on both endothelial cells and migrating CD4+ T cells is upregulated to facilities the motion of the CD4+ in the space between the cells through hemophilic bindings(Wimmer et al., 2019). Trans-cellular migration could constitute one third of the cases of diapedesis(Carman et al., 2007). Interestingly, transcellular migration increases in neuro-inflammatory conditions such as multiple sclerosis(Wu et al., 2016). One of the main differences between transcellular migration and para-cellular migration is the presence membrane fusion process that employs vesicles enriched with caveolae marker (CAV1) as well as vesiculo-vacuolar organelle (VVO) in addition to SNARE proteins that play an important role in transcellular pore formation(Carman & Springer, 2008).

### The origin and the nature of these systems is not yet known

The evolutionary origin of these two systems is not yet known. Whether they constitute the same physiological process is still debated. One argument that supports this notion, is that although trans-endothelial migration is dependent on vesicles formation and SNARE proteins to decrease surface tension(Muller, 2015)/ This observation suggests that these two process are similar and the main difference between them is the leukocytes decision is to probe the space between the endothelial cells or the endothelial cells proper. It has been reported that leukocytes chose the site of entry based on the least resistance principal. Conversely it has been shown that CAV1 expression favors transcellular migration and CAV1-/- mice only exhibit para-cellular migration(Lutz et al., 2017). Similarly in mice muse models of PECAM1-/- mice, lymphocytes migration into the brain exhibited trans-cellular migration(Wimmer et al., 2019). Furthermore in drosophila a para-cellular migration system has been demonstrated(Kunwar, Starz-Gaiano, Bainton, Heberlein, & Lehmann, 2003). However, the system is based in Tre1 protein. However, little is known about Tre1 evolutionary history. Furthermore, the common pathways between para-cellular and trans-migratory pathways are still not well presented.

In this paper we compared the evolutionary origin of each of the components of the two systems using a phylogenetic approach. First, we used an AI text mining method to extract the protein name that are expressed in each migration system. We built phylogenetic trees for each of these components. Our results indicate that both systems contain ancient components such as RhoA which is more than one billion years old. Also both systems share newly divergent components such as ICAM1 and VCAM1. However, there are still several differences between them. One of the major differences is the existence of caveolae in the trans-endothelial group, manifested by CAV1 which exist as ancient as Trichoplax adhaerens and VAMP family that diverged during the emergence of drosophila/ c. elegans. On the other hand PECAM1 that plays a role in para-cellular and not transcellular appeared during the divergence of the vertebrates from FCRL5. Additionally, in drosophila, Tre1 diverged from melatonin GPCRs around the time of divergence of drosophila. Taken together, our results suggest that there are at least three different systems of endothelial migration in the brain. These systems are characterized by several unique components such as PECAM1, CAV1 and TRE1. However, there is considerable interconnection between them.

## Methods

### Material and methods

#### Data used

The motivation of this report was exploring the evolutionary history of para-cellular and trans-cellular trans-migration pathways. In order to do that, we downloaded the human sequences for the proteins known to contribute to either of these two pathways. We employed BLASTP to investigate the similarity between these human protein various species covering 1 billion years, starting with fungi (which one?), common fruit fly (*Drosophila melanogaster*), roundworm (*Caenorhabditis elegans*), sea anemone (*Nematostella vectensis*), Hydra (*Hydra vulgaris*), sea squirt (*Ciona intestinalis*), Lampreys (which type), Zebrafish (*Danio rerio*), Elephant shark (*Callorhinchus milii*), red junglefowl (*Gallus gallus*) and House mouse (*Mus musculus*). In order to increase accuracy we only choose the transcript with the longest number of aminoacids for further analysis. We used the threshold of 1e-10 to accept newly identified proteins as putative candidates. Finally we filtered sequences that do not follow the rest of the specific family domains using CDD.

#### Phylogenetic tree

Phylogenetic analysis was performed in 3 steps. First, investigated proteins investigated were aligned using clustalw function in seaview. After that, we employed ProtTest^27^ to determine the best amino acid replacement model (is that true?). ProtTest outcomes were constructed based on the Akaike information criterion (AIC) that proposed that the best substitution model is LG+I+G+F (is that right?). In our model LG was used as the substitution model, while I is the probability of rate categories given by G (does not make sense). Thirdly, we built our trees by employing PHYML built in in Seaview with five random start for each tree.

#### Ancestral sequence reconstruction

We applied the maximum likelihood method to produce the ancestral sequence of each if the genes investigated. For each gene, we used the ASR algorithm implemented in MEGA6 (Tamura et al., 2013) to build ancestral sequences. This was followed by Blastp against the nearest earlier diverging organism. BLAST outcome was only accepted if the E-value threshold was less than e-10.

#### HHsearch

HHsearch and Needleman-Wunsch methods were used to examine the evolutionary history of each proteins. Only proteins that have already diverged before the inspected protein was considered as a candidate parents (Fredriksson et al., 2003; Nordström et al., 2011). In order to ensure accuracy, the highest HHsearch together with Wilcoxon signed rank were accepted as previously described.

## Results

**Figure 1.**
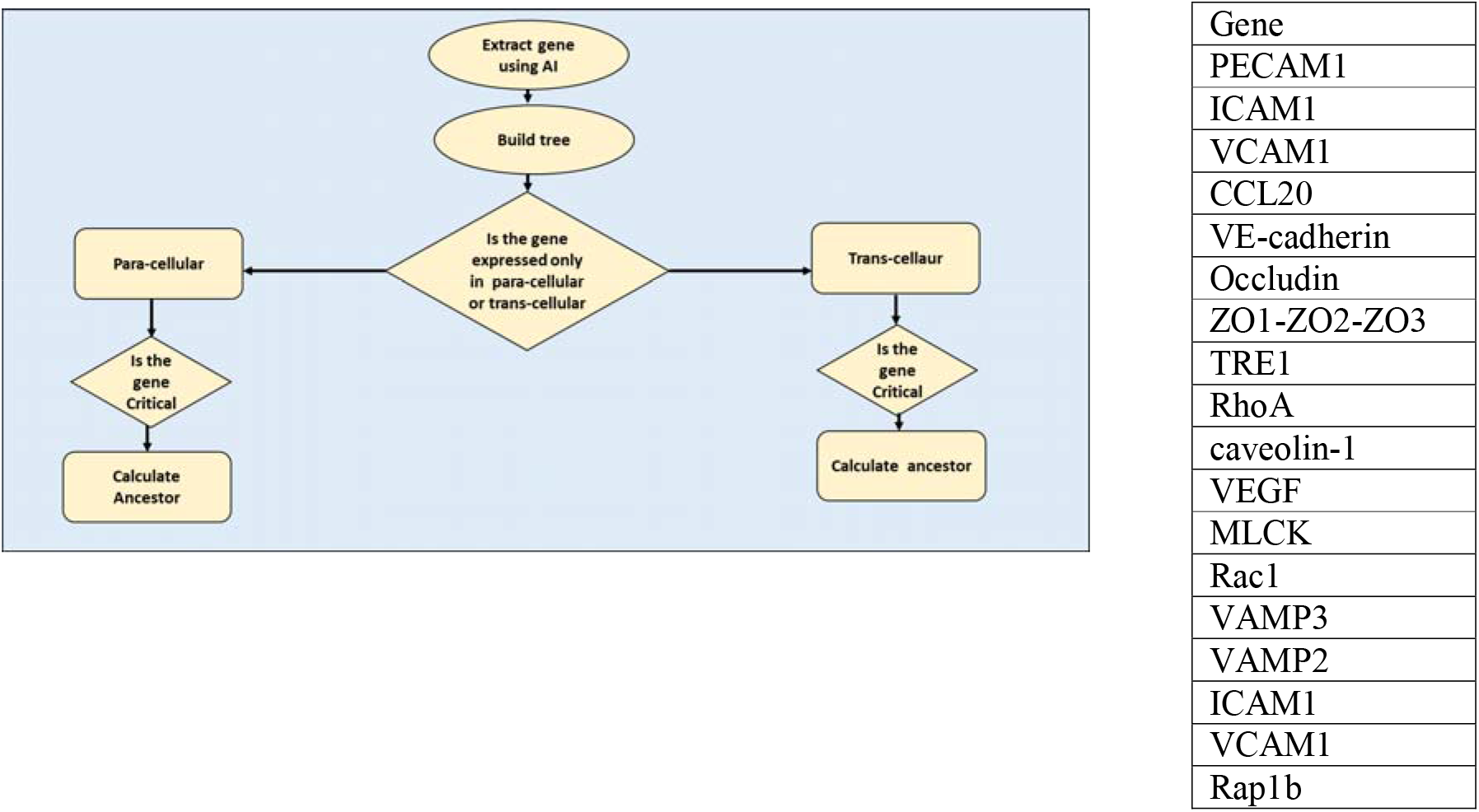
Workflow of this investigation.

### Automatic literature review results (threshold 0.05) on MSE

**Table 1.**
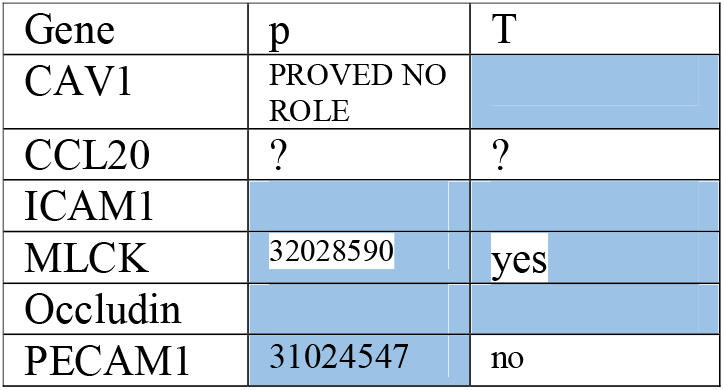
validation of AI literature review

### Phylogenetic analysis of Components of the paraceullar mechanism

We found two PECAM1 genes in Danio, we could not find it in ciona intestinailis, lampreys, elephant shark, *nematostella vectnsis, Hydra vulagris, trichoplax adhaerens* or fungi. Pecam1 is an orphan protein and it was associated with Fcr_like (figure 2a).

**Figure 2.**
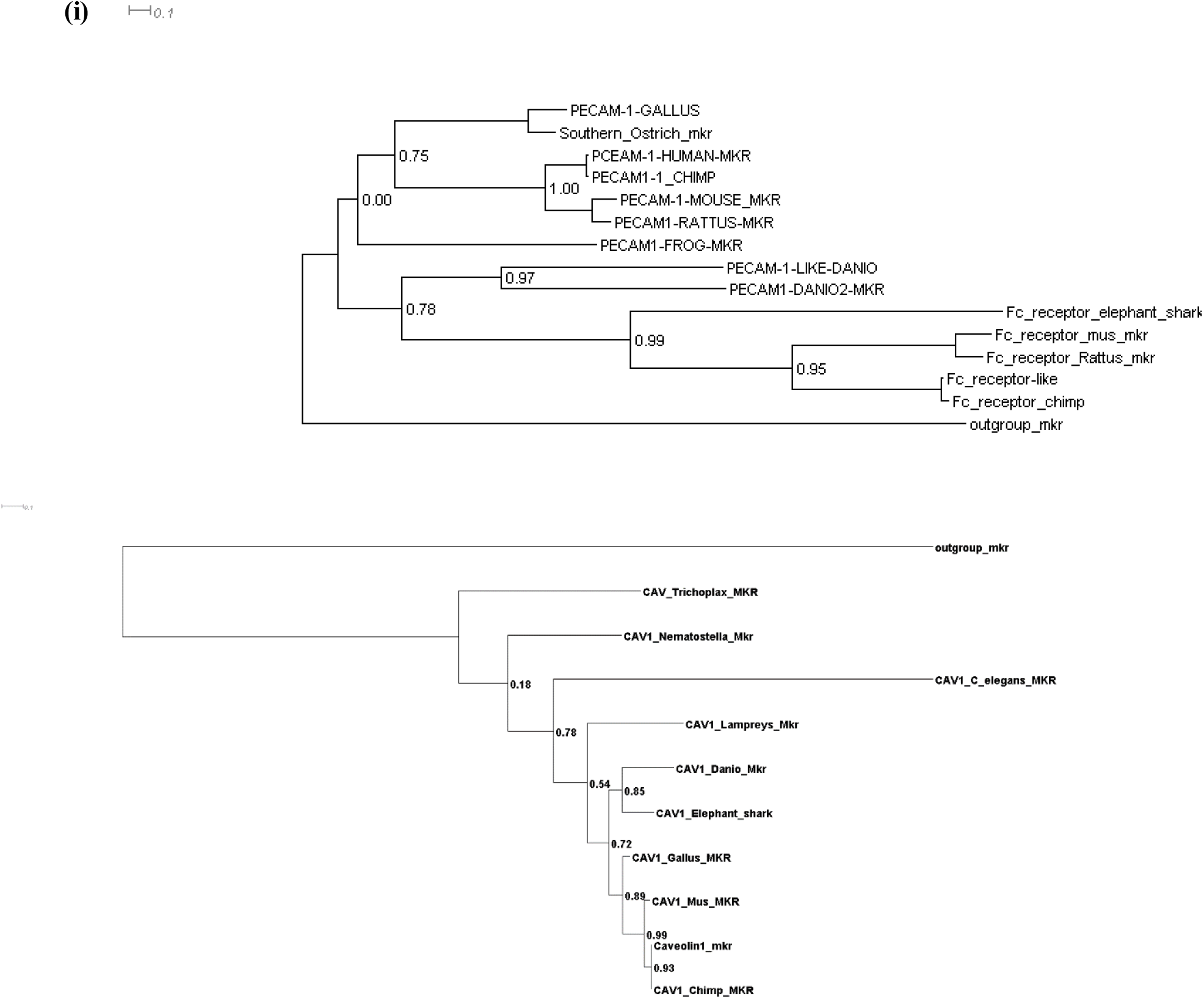
Phylogentic analysis of Components of the trans-ceullar mechanism a)

**Figure 3.**
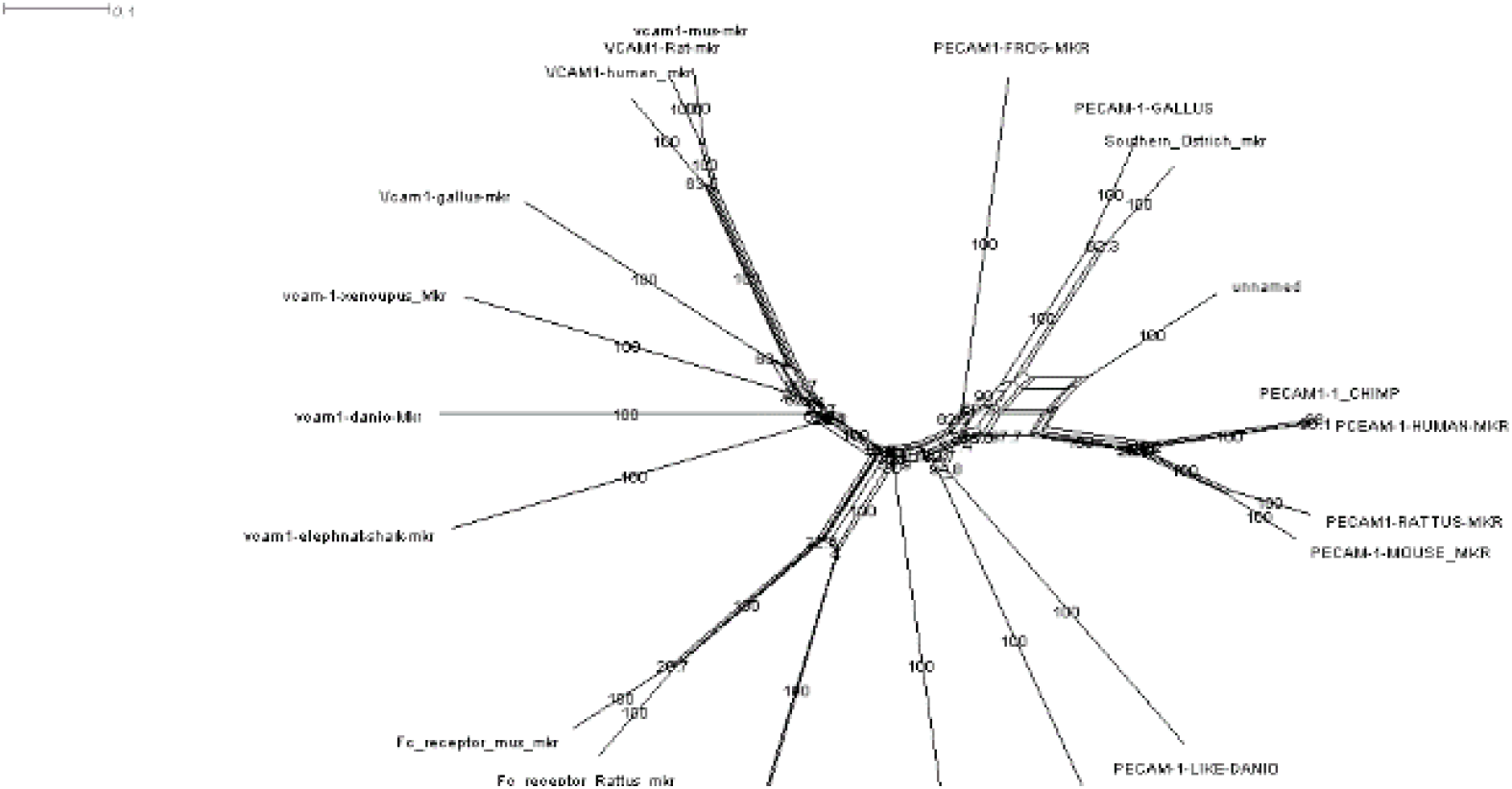
Comparison between the phylogenetic history of the active components of para-cellular and transcellular migration pathways. We investigated the phylogenetic of a) PECAM1 the main active component of para-cellular migration and b) Tre1

**Figure 4.**
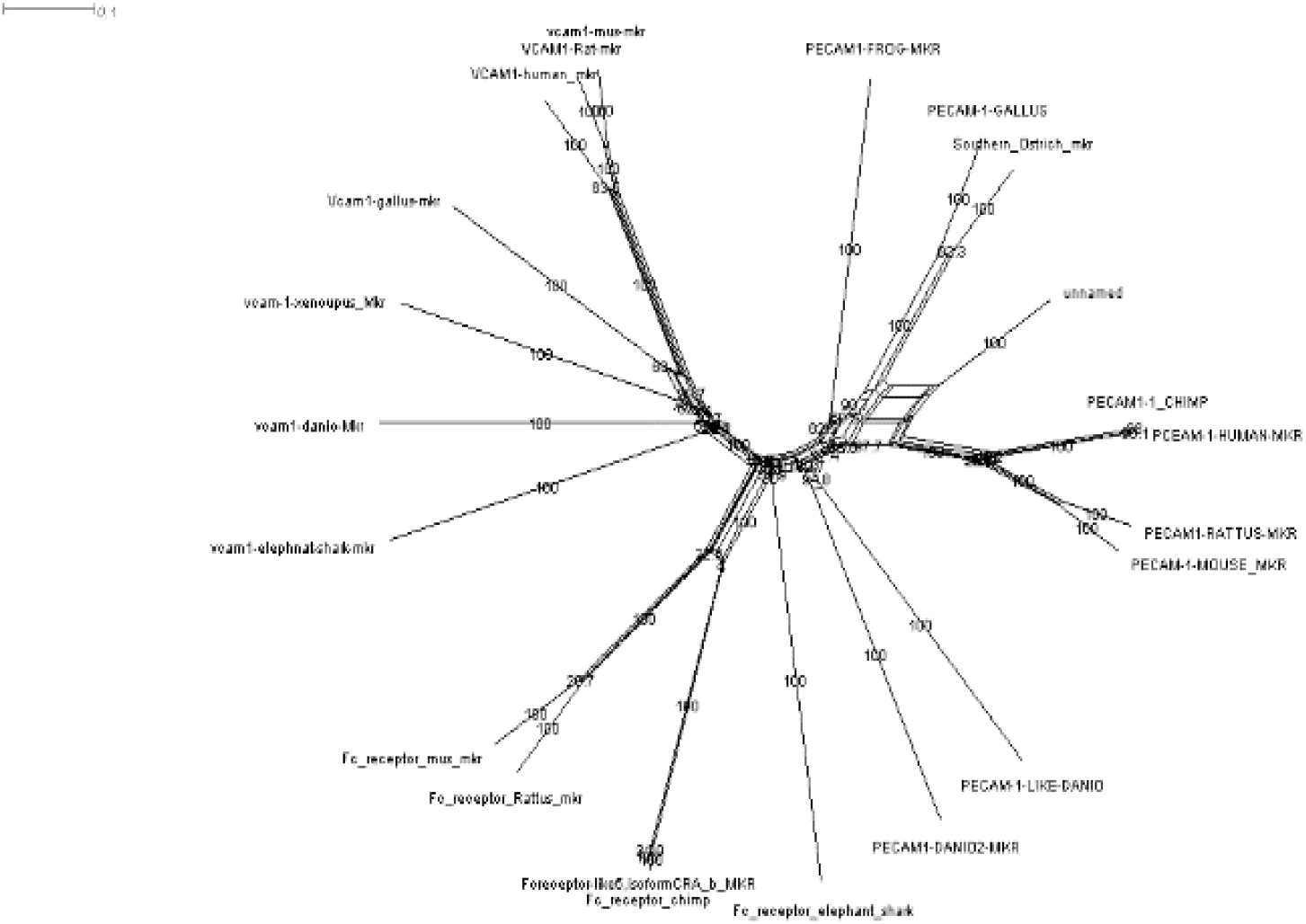
Comparison between the most probable ancestor of the active components of para-cellular and transcellular migration pathways. We investigated the phylogenetic of a) PECAM1 the main active component of para-cellular migration vs.b) CAV1 and c) Tre1

**Figure 5.**
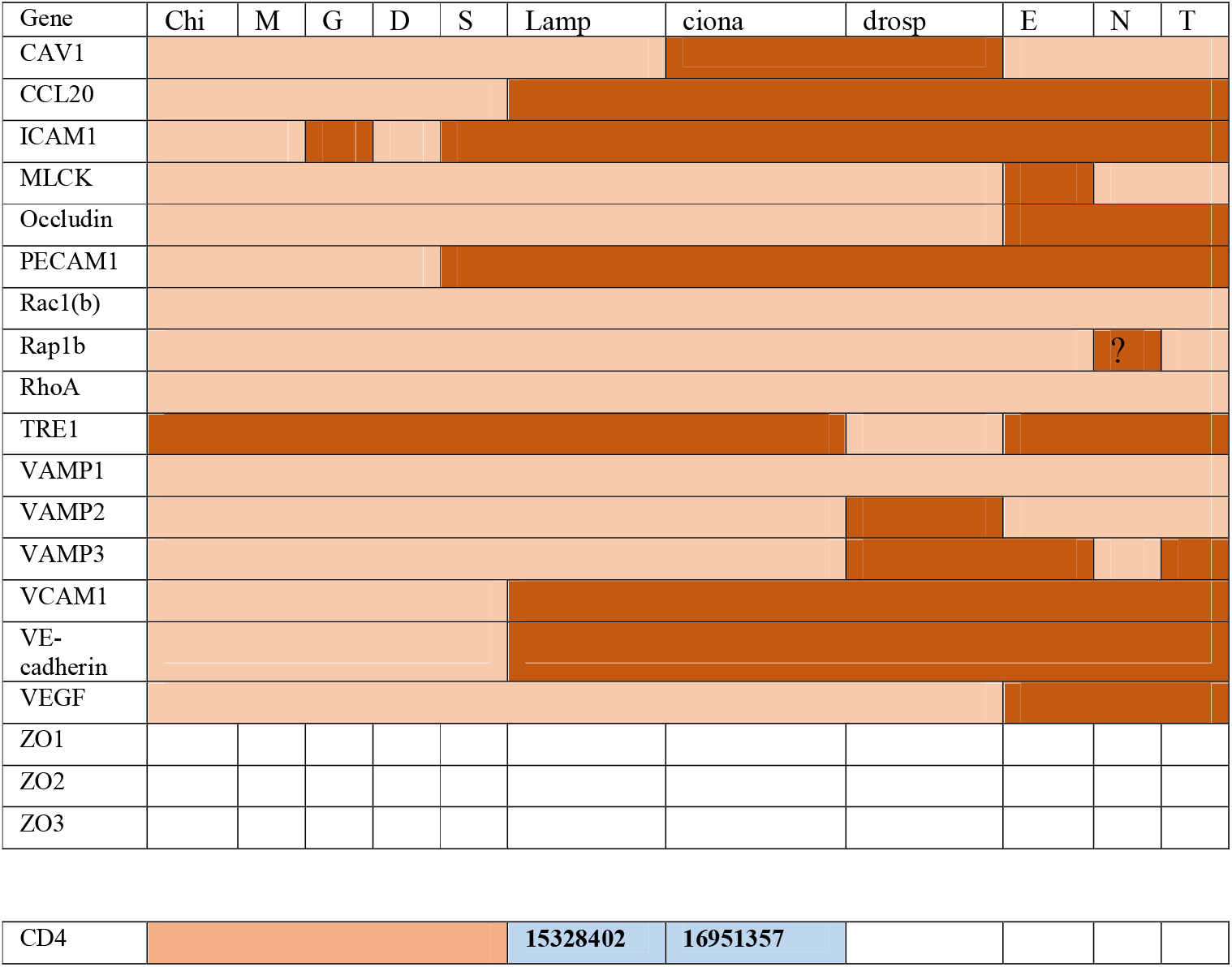
Phylogenetic history of the main components of para-cellular and trans-cellular migration.

## Discussion

In this report we show that that the evolutionary history of transmigration main methods is different. Although the two methods share key mechanisms that help permit transmigration through the endothelial layer of the BBB, they have various components that have different function and different evolutionary history. These key components include CAV1, PECAM and Tre1. CAV1 which is a critical contributor in transcellular migration, has diverged during Trichoplax adhereans and continued to be expressed in major lines except for drosophila and ciona. PECAM1 on the other hand is relatively new as it diverged during the emergence of cartilaginous fish such as elephant shark. Tre1 is more similar to melatonin GPCR, however it only exists in drosophila. Tre1 seems not to be expressed in any species outside insects. Taken together this report indicates that contrary to the ongoing belief of the resemblance in nature and function of paracellular and transcellular migration, they constitute two divergent process.

### Origin of transmigration

#### Two different systems

Although paracellular and transcellular migration process contain various identical components that constitute two divergent evolutionary systems. In mammals both systems are dependent on ICAM1 and VCAM1 interaction with the invading lymphocytes along with the interaction of CCL20 and its receptor CCR7 to initiate the diapedesis process. Our analysis shows that ICAM1 and VCAM1 and CCL20 diverged during the bony fish and cartilaginous fish respectively (figure y). Additionally, both systems use the RhoA pathway to initiate membrane movement. RhoA is an ancient gene which is likely o have diverged before the emergence of Fungi. In paracellular pathway is likely to initiate the phosphorylation of VE-Cadherin, where as this role is not needed in transcellular migration. In Transcellular migration, RhOA initiate the formation of CAV1 in a VAMP mediated fashion, which are also as ancient as Trichoplax. On the other hand, PECAM which diverged during the danio rerio divergence likely from FCR5 form homophilic bonds and initiates the movement of the membrane sideways to increase the possibility of leukocytes infiltration of the brain. Several studies support our findings that these two processes constitute two different process. First it was reported that PECAM1-/- mice favors transcellular migration and not paracellular migration. Secondly, it has been shown that CAV1-/- mice are almost exclusively using transcellular migration. Thirdly it was shown by Muller et al that PECAM1 and CAV1 do not colocalize together in one cellular compartment.to summarize, PECAM1 and CAV1 which are responsible for forming the space for migration in transcellular and paracellular migration receptively are mutually exclusive function-wise, also have different phylogenetic origin and history.

#### Transmigration in lower invertebrates

Trichoplax and nematostella as well as c. elegans constitute a phylogenetic paradox. No immune cells have been reported in them. Trichoplax does not show a rudimentary brain and hence no blood-brain barrier. Nematostella shows two neurons centralization pathways (wnt and BMP). They also express several genes related to astrocytes that play an important role in the BBB. C.elegans have central neurons in a brain-like structure and a BBB like structure which expresses claudin. Interestingly, all three species express CAV1 that is responsible for forming caveolae. Caveolae are not expressed in fungi and in plants, thus it seems to be a metazoan invention. In an interesting study, CAV1 expressed in bacteria initiated the formation of caveolae(Walser et al., 2012). However, these reports have not been yet repeated in any of the lower invertebrates. The question arises about the need for expression of CAV1 in these species. One hypothesis could be that caveolae were initially used to transfer macromolecules. This argument is further supported by other reports that showed that Trichoplax express CAV molecules in order to digest algae(Kaelberer & Bohorquez, 2018). In C. elegans, where a more BBB-like structure exits, it was suggested that a passive diffusion transedothelial of macromolecules is used (Dighe et al., 2019). A system resembling paraceullaur migration is yet to be discovered. Taken together our results suggests that cavloe system existed in earlier vertebrates to help intake of food. Transmigration in drosophila, Regulation of transmigration appeared with the divergence of drosophila & appearance of immune cells as well as in lampreys. What determines the pathway of migration is still an open question and if there is any Cross talk between the two mechanism.

## Acknowledgement

We would like to thank Prof. Macrious Abraham and Prof. Mariam Joachim for their innovative discussions and ideas.

